# A comparison of visual features used by humans and machines to classify wildlife

**DOI:** 10.1101/450189

**Authors:** Zhongqi Miao, Kaitlyn M Gaynor, Jiayun Wang, Ziwei Liu, Oliver Muellerklein, Mohammad Sadegh Norouzzadeh, Rauri C K Bowie, Ran Nathan, Stella X Yu, Wayne M Getz

## Abstract

In our quest to develop more intelligent machines, knowledge of the visual features used by machines to classify objects shall be helpful. The current state of the art in training machines to classify wildlife species from camera-trap data is to employ convolutional neural networks (CNN) encoded within deep learning algorithms. Here we report on results obtained in training a CNN to classify 20 African wildlife species with an overall accuracy of 87.5% from a dataset containing 111,467 images. We then used a gradient-weighted class-activation-mapping (Grad-CAM) procedure to extract the most salient pixels in the final convolution layer. We show that these pixels highlight features in particular images that are in most, but not all, cases similar to those used to train humans to identify these species. Further, we used mutual information methods to identify the neurons in the final convolution layer that consistently respond most strongly across a set of images of one particular species, and we then interpret the features in the image where the strongest responses occur. We also used hierarchical clustering of *feature vectors* (i.e., the state of the final fully-connected layer in the CNN) associated with each image to produce a *visual similarity dendrogram* of identified species. Finally, we evaluated how images that were not part of the training set fell within our dendrogram when these images were one of the 20 species “known” to our CNN in contrast to where they fell when these images were “unknown” to our CNN.

## Introduction

Collecting animal imagery data with motion sensitive cameras is a minimally invasive approach to obtaining relative densities and estimating population trends in animals over time. It enables researchers to study their subjects remotely by counting animals from the collected images^1^. However, due to their complexity, images are not readily analyzable in their raw form and relevant information must be visually extracted. Therefore, human labor is currently the primary means to recognize and count animals in images. This bottleneck impedes the progress of ecological studies that involve image processing. For example, in the Snapshot Serengeti camera-trap project, it took years for experts and citizen scientists to manually label millions of images^2^.

Deep learning methods^3^ have revolutionized our ability to train digital computers to recognize all kinds of objects from imagery data including faces^4, 5^ and wildlife species^2, 6, 7^ (see Appendix 1 for more background information). It may significantly increase the efficiency of associated ecological studies^2, 8^. In our quest to increase the capabilities of machines to communicate with humans, it would be useful to have machines articulate the features they employ to identify objects^9, 10^. This articulation would not only allow machines to converse more intelligently with humans, but may also allow machines to reveal cues that humans are currently not using for object identification, which could then make humans more effective at such identification tasks. Before we can do this, however, we must identify the human-coherent, visual features used by machines to classify objects. To the best of our knowledge, none of the few existing studies that use deep learning for animal classification concentrate on this issue. As such, they lack the necessary transparency for effective implementation and reproducibility of deep learning methods in wildlife ecology and conservation biology.

To identify such features in the context of classification of wildlife from camera trap data, we trained a Convolutional Neural Network (CNN)^7, 11^ using a deep-learning algorithm (VGG-16: as described in^12^ and in Appendix 4) on a fully annotated dataset from Gorongosa National Park, Mozambique (Figure 5) that has not previously been subjected to machine learning. After training, we interrogated our network to better understand the features it used to make identifications by deconstructing the features on the following three aspects of our implementation: 1) localized visual feature, 2) common intraspecific visual features, and 3) interspecific visual similarities (Figure 8).

We used Guided Grad-CAM (GG-CAM) methods—a combination of Guided Back-propagation (GBP)^13^ and gradient-weighted class activation mapping (Grad-CAM)^14^)—on the last convolutional layer of our trained network to extract localized visual features of single images and compare the results with visual descriptors used by human classifiers to identify species in our image set. We found that the features used by the CNN to identify animals were similar to those used by the human. Next, we used the Mutual Information (MI) method^15, 16^ to generalize within-species features as an extension of the localized visual feature of single images. Finally, we used hierarchical clustering^17^ on the CNN feature vectors to further inspect the visual similarities between animals species learned by the CNN. We found that the relative visual similarities emerged during training process were similar to human knowledge as well. We also used an experimental testing dataset that have 10 “unknown” animal species to measure the relative familiarity of species to the CNN. The results implied that visual similarities could be used to identify visually distinct “unknown” animal species. In the Discussion section, we show a brief example of how interpretations of CNNs can help to understand the causes of misclassification and to make potential improvements of the method.

## Methods and results

### 0.1 Model training

Before interpreting a CNN, we firstly trained a VGG-16^12^ on a fully annotated dataset from Gorongosa National Park, Mozambique (Figure 5). To increase the convergence fidelity of our learning algorithm in extracting species-specific visual features, we confined our training images to only the 20 most abundant species (ranging from 473 images of hartebeest to 28,008 images of baboons, Figure 6). Adding some of the rarer species would have degraded the overall performance of the network because the network has fewer images to use in extracting species-specific visual features or species-specific combinations from a more general set of visual features that are not on their own species-specific.

Under this somewhat ad-hoc constraint on the number of species, after pruning out all images not containing the 20 most abundant species, we split the remaining 111,467 images at random into training (85% of images), validation (5% of images; for tuning hyperparameters listed in Table 1), and testing (10% of images; for evaluating accuracy) subsets. We used a deep learning algorithm (VGG-16)^12^ (see Appendix 3 for implementation details), which we then evaluated for accuracy once trained (Figure 7; overall accuracy was 87.5%, and average accuracy across the 20 species was 83.0%, ranging from a high of 95.2% for civet to 54.3% for Reedbuck—see Figure 7(a). Data distribution, habitat types or time of day when the image was captured, did not affect the performance of the deep learning algorithm.)

### Localized visual feature and human comparison

Then, we used GG-CAM methods, which combines the output from GBP^13^ and Grad-CAM^14^, on the last convolutional layer of our trained network, where feature localization occurs (see Appendix 4). We note that Grad-CAM captures the most discriminative image patch, GPB captures visual features both within and outside of the focal Grad-CAM patch, and GG-CAM captures the features most salient to the actual discrimination process (Figure 9). We then inspected the GG-CAM images produced by our CNN relative to the original images in order to assess what sort of localized visual discriminative features were being extracted from the original images (Figure 1); in this manner, we obtained information on the inner mechanism of deep learning classification^18, 19^.

**Figure 1.**
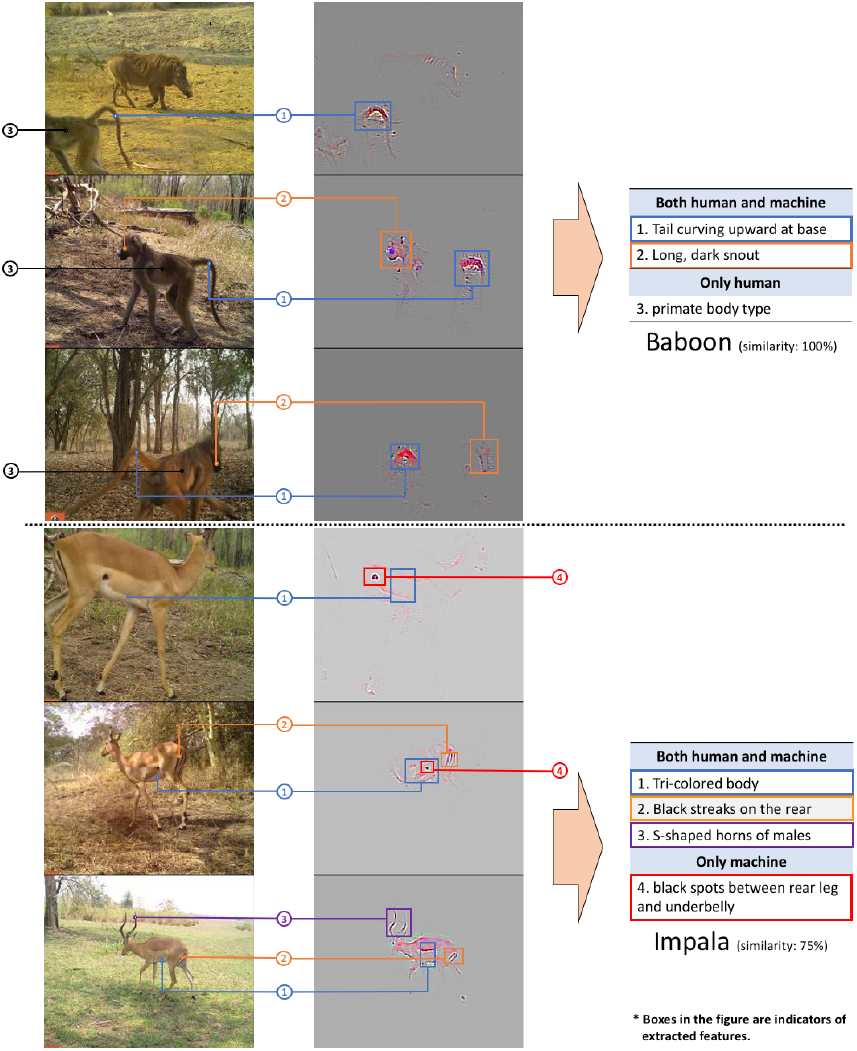
GG-CAM generated localized discriminative visual features of randomly selected images of baboon and impala. For classifying baboons, the CNN focuses on faces and tails. For impalas, the CNN uses the contrast between the white underbelly and dark back, black streaks on the rear, and black spots between the rear legs and underbelly. Most of the features extracted by the CNN have counterparts (similar focal visual components) in the human visual descriptors (indicated by the colors and agreed upon by at least 2 of 4 authors). The similarity in descriptors is calculated as the percentage of extracted features that are similar to the corresponding human descriptors (further detail in Figure 10).

We compared the features extracted by GG-Cam to the visual descriptors used by human classifiers to identify species in our image set (as described in Appendix 2 and Table 4). We calculated the relative similarity for each species, defined as the percentage of extracted features that were similar to those regularly used by humans. The similarity was agreed upon by at least two of four authors (ZM, KMG, ZL and MSN) who scored nine randomly selected images for each species. The trained CNN uses features similar to those used by humans to identify most of the animal species (Figure 10). The mean similarity across species was 0.71 (standard deviation: 0.18). Figure 1/Baboon shows that our CNN uses faces and tails to identify Baboon images. Both of the two features have counterparts (similar focusing areas) in Table 4/Baboon. In Figure 1/Impala, besides the black streaks on the back ends, the line separating the colors of the upper body from the white underbelly and S-shaped horns, the CNN also appears to consider the black spots between the rear legs and bellies of impala as a discriminative feature. This feature, although not included in the most-used descriptors, is a good example of a discriminatory feature traditionally overlooked by humans but now identified by our CNN as salient for use in future identifications.

### Common within-species features

Next, we used the Mutual Information (MI) method^15, 16^ to extend the features of single images to within-species features of each animal species. We calculated the MI scores for each of the neurons in the last convolutional layer of our CNN to indicate their importance to all images of one of the selected species (Appendix 4). In short, for each of these neurons, we obtained 20 species-specific MI scores. For each species, we identified the five neurons in the last convolutional layer that produced the five highest scores. We then identified the “hottest” 60x60 pixel patch (within-species features) to which each of these top five neurons responded in each image (e.g. Figure 2). These features generalize across all images within the same species, as illustrated in Figure 11. Most results are associated with distinguishable visual features of the animals, for example, black spots on civets, an elephant trunk, quills on porcupines, and white stripes on nyala. However, visual similarities of animal species are not the only information our CNN uses to identify species. CNNs also use information such as the presence of trees in the background to identify species frequenting woodlands, especially when most of the images are from similar environments or the same camera-trap locals (e.g., image patches of the top1 neurons of wildebeest and porcupine in Figure 11).

**Figure 2.**
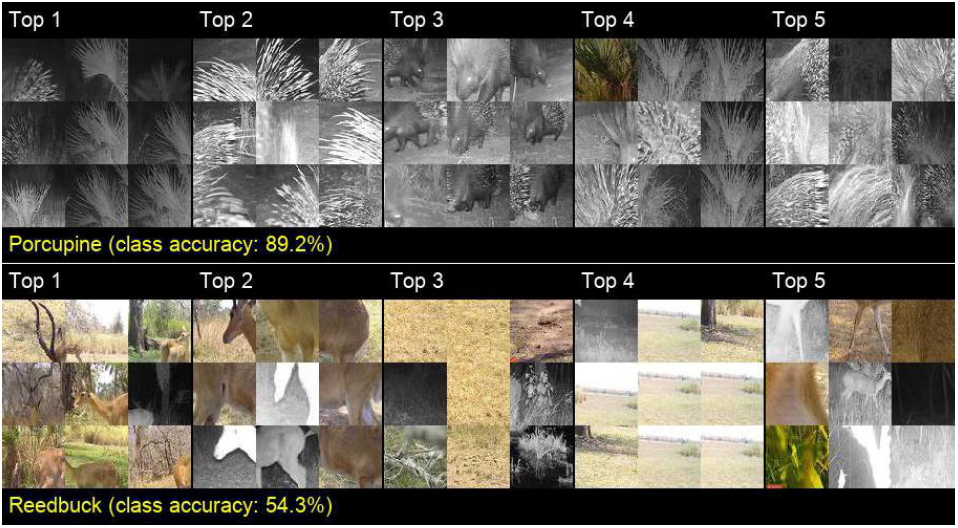
Image patches that respond most strongly to the five neurons with the highest MI scores of porcupine and reedbuck. The leftmost set of nine 60x60-pixel patches are extracted from nine camera-trap images that include a species of interest. These nine images are selected at random from training images of the said species. In each of the nine cases, the extracted patches are centered around the “hottest” pixel (i.e., highest response) of the neuron (in the last convolutional layer of our CNN) that has the highest MI score (Appendix 4) for the said species class. The remaining four sets of nine patches are equivalently extracted for the neurons with the next four highest MI scores. These patches provide a sense of the within-species features to which the neuron in question responds. The higher the class accuracy, the more closely correlated these image patches are for the species of interest. For example, in the relatively accurately identified porcupine set (89.2% accuracy), the first neuron (Top 1, of the upper set) responds to palm plants that appear in most of the training images that also contain porcupines. The second neuron (Top 2) responds to the quills, while the third neuron (Top 3) responds most strongly to bodies with faces. On the other hand, in a much less accurately identified reedbuck set, the first neuron (Top 1, of the lower set) appears to respond to branch-like structures, including tree limbs and horns, but the patterns are less consistent than for the porcupine. Note that some sets of patches are primarily backgrounds (e.g., Top 1 upper set and Top 4 lower set), from which we can infer that our CNN learns to associate certain backgrounds with particular species. Such associations, however, only arise because particular cameras produce common backgrounds for all their images, thereby setting up a potential for a camera-background/species correlation that could well disappear if additional cameras are used to capture images. Similar sets of images are illustrated for other species in Figure 11.

### Interspecific visual similarities

Finally, we generated a visual similarity dendrogram for all species by applying hierarchical clustering^17^ to the CNN feature vectors of 6000 randomly selected training images, i.e., the outputs of the last fully-connected layer (which is of dimension 4096 in Euclidean space) of our trained CNN (see Appdendix 0.1). This dendrogram (Figure 3) is an abstract representation of how images of species are separated in the feature vector space. It also provides a means for quantifying how visually similar the 20 animal species are to our trained CNN. Similar animals are measurably closer together than those that are visually distinct (e.g., striped versus spotted; long-tailed versus no-tail), irrespective of their phylogenetic distance. Thus, though most of the antelopes are grouped together (from sable to reedbuck), the large bull-like herbivores (wildebeest and buffalo) and pig-like mammals (warthog, porcupine, and bushpig) are also grouped together even though they may belong to different families or orders (Figure 3). A well-learned feature vector space can also help identify images that differ in some way from those on which the CNN has been trained^20, 21^.

**Figure 3.**
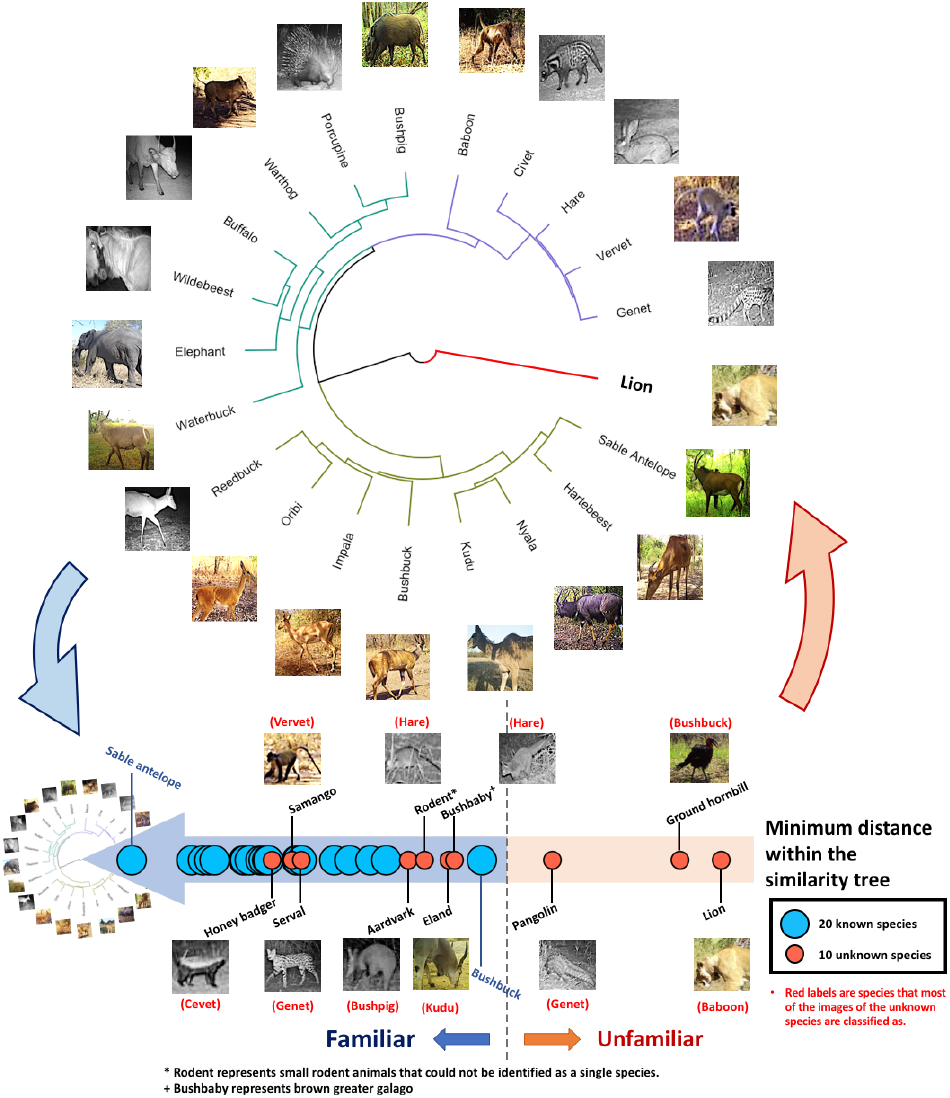
Visual similarity tree for our trained CNN and relative familiarity of 30 species to the CNN. The similarity tree is based on hierarchical clustering of the response of the last fully-connected layer in our trained CNN to 6000 randomly selected training images of particular species (i.e., feature vectors of the images). The leaves represent feature vector centroids of 300 training images of each species, and their relative positions in the tree indicate the Euclidean distances between these centroids in the feature space. In the similarity tree, the more similar the response of this layer to two species, the more tightly coupled they are in the tree. Green, purple, and brown branches correspond to three primary clusters that appear to be a small to medium sized antelope cluster, an animals-with-prominent-tail or big-ears cluster (though baboons seem to be an outlier in this group), and a relatively large body-to-appendages group (with waterbuck the outlier in this group). Also indicated are the relative familiarity of 30 species (including 10 unknown species) to the CNN. To measure the relative familiarity, we used an experimental testing dataset of 30 animals species to calculate the minimum distances of each 30 species within the tree (i.e., we treated each of the 30 species as a new species, either known or unknown, generated a new similarity tree, and calculated the Euclidean distances between these species and their “closest” species in the feature space, Appendix 4). This experimental testing dataset contains 20 randomly selected images that are not used during training of 30 animals species (with 20 known species and 10 unknown species). Although our CNN was not trained to identify unknown animal species, some animal species can still be identified as “unfamiliar” by their distances within the similarity tree. The blue circles represent the 20 known species and the orange circles represent the 10 unknown species. Most of the known species are clustered on the left side, which indicates that the testing images of known species are relatively close to their corresponding feature vector centroids. On the other hand, three of the unknown animals are at larger distances within the similarity tree. When places in the similarity tree (e.g., the red branch of lion), the feature vectors of those animal species can differ greatly from those of the known species. However, except for pangolin, ground hornbill, and lion, the other seven unknown animal species fit closely within the similarity tree, which indicates that the CNN did not learn features that separate the known species from the morphologically close unknown species.

We used an experimental testing dataset, which contained 20 randomly selected images each of 20 known species (species used during training) and 10 unknown animals (Appendix 4), to measure the relative familiarity of both known and unknown species to the CNN. The relative familiarity was calculated as the Euclidean distances between the feature vector centroids of both known and unknown species in the experimental testing dataset to the centroids of training data that constructed the similarity tree (Figure 3, also see Appendix 4). The known species were always relatively close to the 20 feature vector centroids of the training data, whereas some of the unknown species (pangolin, ground hornbill, and lion) were measurably far from the 20 known species; most of the unknown species were still relatively close to the known species, indicating that those animal may share features with the 20 known species (e.g. black spots are shared by civets and servals)^22^.

## Discussion

Understanding the mechanisms of deep learning classifications of camera-trap images can help ecologists determine the possible reasons for misclassification and develop intuitions about deep learning, which is necessary for method refinement and further implementation. For example, Figure 7(a) indicates that reedbuck is the least accurately classified species by the CNN. The confusion matrix^23^ of testing results (Table 3) reveals that many reedbuck images are classified as oribi (8%), impala (12%), and bushbuck (12%). Figure 3 shows that reedbuck is close to oribi, impala, and bushbuck in the feature vector space learned by the CNN, which partly explains misclassification. Further, by examining the localized visual features of the misclassified images, we can gain a clearer sense of the reasons for misclassification. Figure 4 depicts examples of misclassified reedbuck images. Although the CNN can locate the animals in most of the images, it is challenging for the CNN to classify the images correctly when the distinct features of the species are obscured or multiple species are in the same scenes.

**Figure 4.**
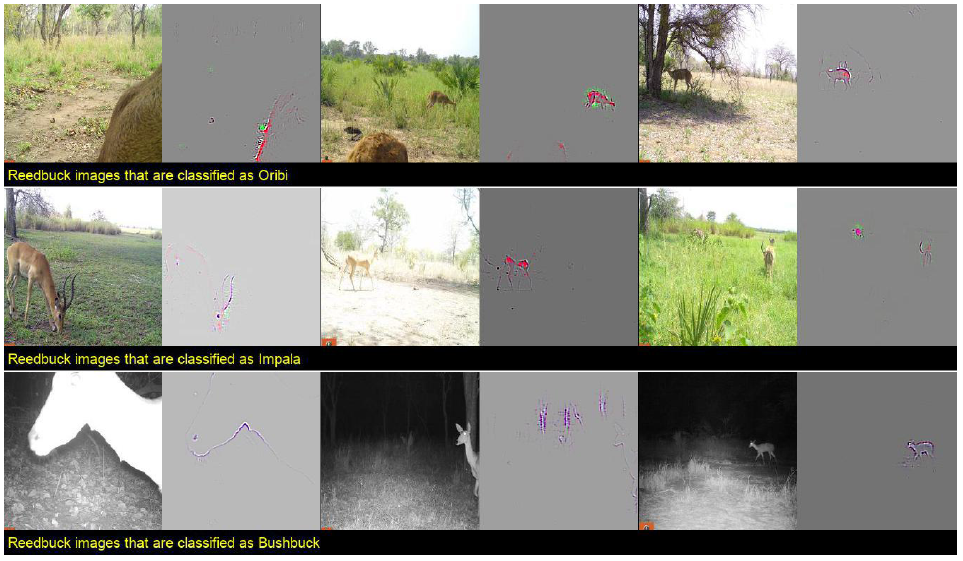
Examples of reedbuck images that are misclassified as oribi, impala, and bushbuck, with corresponding localized discriminative visual features. Although the CNN can locate animals in most images, it is hard for the machine to find distinct features from: 1) images with animals that are far away in the scene; 2) over-exposed images; 3) images that capture only parts of the animal; and 4) images with multiple animal species. In many of these cases, the other species are indeed present in the scenes, and are often in the foreground. This problem is an artifact of the current labeling process and remains to be resolved in the future. For example, the animal in the leftmost image on the second row that is classified as impala is an impala. The CNN correctly classifies this image based on the animal. However, this image was also labeled as reedbuck because the extremely small black spots far in the background are reedbuck. When two species appear in the same scene, the same image is saved twice in the dataset with different labels corresponding to different species in the scene. This labeling protocol can confuse the CNN and remains a problem that must to be resolved in the future.

Deep learning has become a core component of data science and fields using big data. Ecology has been no exception, with its shift towards the machine learning methods in ecoinformatics^24, 25^, including problems in conservation biology^26^, as well as the merging of data analytics with the scientific method^27^. This shift requires that new methods, including models from machine learning and artificial intelligence, are accessible and usable by ecologists^28^. Our paper provides practical steps in model interpretation to help ecologists take advantage of deep learning as a cutting-edge approach for future research and for overcoming major methodological roadblocks. The interpretations described in this paper are steps toward a more informed use of deep learning methods. Future research involving the training of CNNs to identify individuals in ecological studies, whether for purposes of species classification, conservation biology, sustainability management, or identification of specific individuals in their own right^29, 30^ (e.g., in behavioral studies), can follow the methods presented here to identify the sets of features being used to classify individuals. This information may then be used in creative ways yet to be imagined to improve CNN training and, hence, raise the level of performance of CNNs as an aid to analyzing ecological data.

## Acknowledgements (not compulsory)

Thanks to T. Gu, A. Ke, H. Rosen, A. Wu, C. Jurgensen, E. Lai, M. Levy, and E. Silverberg for annotating the images used in this study, and to everyone else involved in this project. Data collection was supported by J. Brashares and through grants to KMG from the NSF-GRFP, the Rufford Foundation, Idea Wild, the Explorers Club, and the UC Berkeley Center for African Studies. We are grateful for the support of Gorongosa National Park, especially M. Stalmans in permitting and facilitating this research. ZM was funded in part by NSF EEID Grant 1617982 to WMG, RCKB and RN, and was also supported in part by BSF Grant 2015904 to RN and WMG. Thanks to Z. Beba, T. Easter, P. Hammond, Z. Melvin, L. Reiswig, and N. Schramm for participating in the feature survey.

## Author contributions statement

This study was conceived by ZM, JW, ZL, KMG, AM and OM. Code was written by ZM and JW and the computations were undertaken by ZM with help from JW, ZL and SXY. The main text was drafted by ZM and WMG with contributions, editing and comments from all authors, particularly RN and RCKB. The appendicies were primarily written by ZM, JW, ZL and KMG. KMG collected all data, oversaw annotation, and conducted the survey. ZM created all figures and tables in consultation with WMG, ZL and SXY.

## Appendix 1: Background

### Camera trap studies

Camera traps have become an increasingly popular tool for the remote monitoring of wildlife populations^31^. They are a low-cost method for gathering data on an entire wildlife community, and are especially useful for studying rare, cryptic, or nocturnal species, including many species of conservation concern^32^. Long-term camera trap datasets can be useful for monitoring trends in populations over time, and novel analyses enable the estimation of relative densities and abundances across time and space^33^. Camera traps set across spatial gradients of environmental heterogeneity can also be used to understand environmental and anthropogenic drivers of wildlife distributions. Analyses can range from simple statistical tests (e.g., Analysis of Variance of relative activity across habitat types) to complex models (e.g., Bayesian hierarchical occupancy models that account for imperfect detection and incorporate multiple predictors of occupancy and detection^34, 35^;). Finally, images or videos from camera traps provide insight into animal behavior, including movement and migration patterns, foraging and anti-predator vigilance, or reactions to experimental stimuli^1^. However, in all of these cases, camera trap studies are limited by the inefficiencies of data processing and the manual classification of species and behaviors in images. Deep learning with CNNs has proven to drastically improve the efficiency of relevant studies. In this paper we demonstrate the mechanisms of deep learning in detail.

### Deep learning in camera-trap classification

Deep learning is a subdomain of machine learning that uses algorithms inspired by biological neural networks^3^. It has gained much attention among ecologists in recent years^36^, with animal species identification from camera trap images using CNNs being one of the most popular applications^2, 7, 8, 37–39^. Chen et. al.^37^ made the first attempt to automatically classify camera trap images with deep learning methods. They achieved only 38% classification accuracy on their 20,000-image dataset, and suggested that, with enough training data, deep learning can surpass other existing methods. Gomez et. al.^8^ harnessed deep learning with transfer learning, a method of fine-tuning, to identify animal species in the Snapshot Serengeti dataset^40, 41^ and achieved over 80% classification accuracy using large amounts of data. Further, Norouzzadeh et. al.^2^ trained multiple CNN architectures on the same dataset as Gomez et. al. and achieved a classification accuracy in excess of 95%, the current state-of-the-art performance for deep learning models in camera-trap studies. However, to our knowledge, there are no studies specifically explaining the mechanisms of deep learning that facilitate classification of animals with such a high degree of accuracy. In addition, the only big camera-trap dataset to which this method has been applied is the Snapshot Serengeti dataset. In this paper, we implement deep learning on a dataset that has not been studied previously and illustrate three approaches to interpretation to reveal the mechanisms of CNNs qualitatively.

### Basic mechanisms of CNNs

Convolutional neural networks (CNNs) are one of the most frequently used deep networks in computer vision From AlexNet^42^ to VGG^12^ and ResNet^43^, the capacity of modern CNN architectures has advanced rapidly, resulting in high recognition accuracies that make abundant real-world applications possible. Modern CNN architectures typically have three types of layers – convolutional layers, pooling layers and fully-connected layers – which gradually transform an input image into a predicted category label. For instance, the VGG-16 network (architecture used in this paper) has 13 convolutional layers, 5 pooling layers, and 3 fully-connected layers; it takes a 227 *×* 227 image as input and predicts a 1-of-1000 category label as output. Convolutional layers in CNNs consist of local filters or neurons and are designed to capture spatially-distributed local traits such as edges, parts and textures^44^. Pooling layers account for the larger receptive field of the deeper convolutional layers, i.e. the subsequent convolutional layers assemble the previously learned local traits into more globally-perceived shapes and configurations^18^. Fully-connected layers abstract all of the local and global traits into high-level semantic concepts like categories and attributes^5^. All the parameters in the CNNs are learned by minimizing the errors between prediction and ground-truthed data through a layer-by-layer updating process called back-propagation. In this work, we interpret the inner representations of CNNs qualitatively and quantitatively by examining the relationship between neurons and ecological data.

## Appendix 2: Data

### Study site

The camera-trap data comes from a long-term research program in Gorongosa National Park, Mozambique (18.8154°S, 34.4963°E) (Figure 5). The dataset used in this analysis was collected from June to November of 2016. The goal of this program is to examine the spatial distribution of large mammal species in the park and to monitor the restoration of the park’s wildlife following decades of civil war. The 3,700 km^2^ park encompasses a range of habitats, including a mix of grassland, open woodland, and closed forest. KMG placed 60 motion-activated Bushnell TrophyCam and Essential E2 cameras in a 300 km^2^ area in the southern area of the park. Each camera was mounted on a tree within 100 meters of the center of a 5 km^2^ hexagonal grid cell, facing an animal trail or open area with signs of animal activity. To minimize false triggers, cameras were set in shaded, south-facing sites that were clear of tall grass. Cameras were set to take 2 photographs per detection with an interval of 30 seconds between photograph bursts.

**Figure 5.**
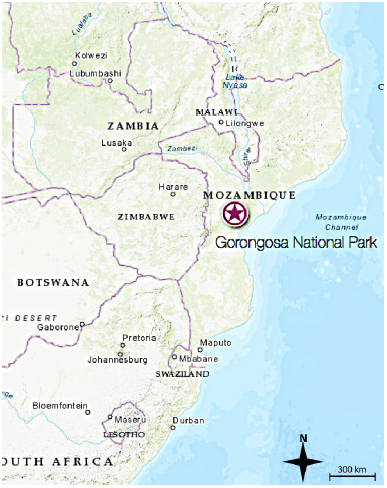
Location of Gorongosa National Park, Mozambique

### Human classification

The species in all images were classified and manually annotated independently by two different researchers trained on a list of example images and corresponding visual descriptors of each species; this list was created by KMG before the manual annotation and was iteratively updated as the annotation progressed. All classifications were confirmed by KMG prior to this project.

We conducted a survey to determine the features that were regularly used by humans to identify each of the 20 species in this study. For each species, respondents were asked to select features that they regularly look for and/or use as clear diagnostic features that identify the species. We provided respondents with KMG’s list of all visual descriptors used in training materials, and included an option of adding additional descriptors not mentioned. The survey had 13 respondents, all of whom have extensive experience classifying camera trap images from Gorongosa National Park, including those used in this study. We considered a feature to be regularly used by humans if at least 5 of the 13 respondents selected it (Table 4).

### Data description

The dataset contains a total of 30 animal species. In this paper, we use data from the 20 most commonly photographed mammal species during data collection for higher training performance and more accurate feature extraction (Figure 6) and omit rare species with less than 350 images, as well as images that are annotated as empty (Figure 6) during training and performance testing. The 20 species include: African buffalo (*Syncerus caffer*); African elephant (*Loxodonta africana*); African savanna hare (*Lepus microtis*); baboon (*Papio cynocephalus*); blue wildebeest (*Connochaetes taurinus*); bushpig (*Potamochoerus larvatus*); Cape bushbuck (*Tragelaphus sylvaticus*); civet (*Civettictis civetta*); southern reedbuck (*Redunca arundinum*); crested porcupine (*Hystrix cristata*); greater kudu (*Tragelaphus strepciseros*); impala (*Aepyceros melampus*); large-spotted genet (*Genetta tigrina*); Lichtenstein’s hartebeest (*Alcelaphus buselaphus*); nyala (*Tragelaphus angasii*); oribi (*Ourebia ourebi*); sable antelope (*Hippotragus niger*); vervet monkey (*Chlorocebus pygerythrus*); warthog (*Phacochoerus africanus*); and waterbuck (*Kobus ellipsiprymnus*). Besides the labels, each image has information on camera shooting times and animal habitat. There are seven types of habitat in the dataset: sparse woodland; sparse to open woodland; open woodland; open to closed woodland; closed woodland; closed woodland to forest; and forest.

When inspecting the relationship of known and unknown animal species and the similarity tree (i.e. 20 mean feature vectors), we created an experimental testing datasets with 20 randomly selected images of each of the 20 species used during training and the 10 excluded species. These 10 species are: aardvark (*Orycteropus afer*); bushbaby / brown greater galago (*Otolemur crassicaudatus*); eland (*Taurotragus oryx*); honey badger (*Mellivora capensis*); lion (*Panthera leo*); samango (*Cercopithecus albogularis*); serval (*Leptailurus serval*); southern ground hornbill (*Bucorvus leadbeateri*); Temminck’s ground pangolin (*Smutsia temminckii*); and rodent (multiple rodent species).

**Figure 6.**
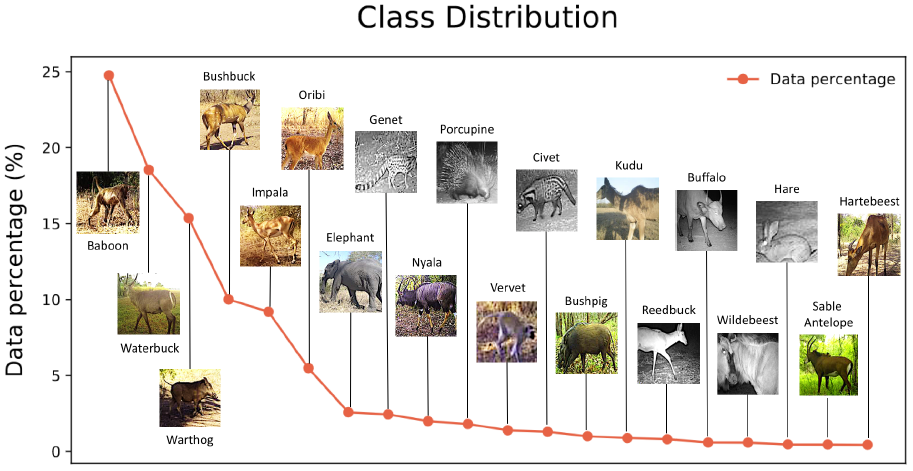
Distribution of 20 different animal species in the 111,467 images used to train, validate and test (85%, 5% and 10% respective split) our CCN. More than 60% of the images include the first three species. The overall accuracy of CNN was 87.5% and average accuracy across the 20 samples was of 83.0% (range was Civet 95.2% - Reedbuck 54.3%, as detailed in Fig 7a)

### Data preprocessing

We first grouped the images by camera shooting events. At each shooting event, when the motion sensors detect motion, the cameras captured two sequential images within one second. Image pairs of the same shooting events often are similar in appearance, and the training performance of the model can be underestimated if images from the same image pair are separated into training and test sets. Thus, we maintained the image pairs in the analysis in order to prevent a negative bias of the CNN learning process. We then randomly split the image groups into training, validation and testing sets with 85%, 5% and 10% of the datasets. In this section, we provide details and results of the implementation of the model along with the results of three methods of interpretation: 1) localized visual features; 2) common within-species visual features; and 3) interspecific visual similarities.

## Appendix 3: Training details Model implementation

We trained a VGG-16^12^ CNN architecture to classify camera-trap images with class-aware sampling^45^. The output of the CNN classifier is a 20-dimensional vector, with each dimension representing the classification probability for an animal species (classification score). The use of class-aware sampling helps to improve classification accuracy for unbalanced datasets.

We made use of PyTorch^46^, a deep learning framework, to implement and train the CNN. The weights were initialized from an ImageNet^47^ pretrained model. The initial learning rate was 0.01, which decreased every 15 epochs. The best model was obtained at epoch 40 where the classification accuracy on the validation dataset was the highest. The loss function used to train the CNN was Softmax cross-entropy loss. All the input images for training were firstly downsized to 256 *×* 256, then were randomly cropped to 224 *×* 224 with a random horizontal flip at rate 0.5. Values of the hyperparameters used for training are listed in Table 1.

**Table 1.** Hyperparameters

| Parameters | Values |
| --- | --- |
| Input image size: | $256 \times 256$ |
| Random crop size: | $224 \times 224$ |
| Random horizontal flip rate: | 0.5 |
| Batch size: | 256 |
| Training epoch: | 40 |
| Initial learning rate: | 0.01 |
| Momentum: | 0.9 |
| Learning rate reduce at: | every 15 epochs |
| Learning rate reduce by: | 0.1 |
| Regularization: | None |

### Assessing accuracy

The classification accuracies of the model are measured by both micro-averaged accuracy (over‐ all accuracy) and macro-averaged accuracy (averaged accuracy per class) in the testing set (Table 2). Figure 7 depicts classification accuracy by animal species, habitat, and camera shooting time of testing images.

**Table 2.** Testing accuracy

| Metric | Accuracy |
| --- | --- |
| Overall micro accuracy: | 87.5 % |
| Overall macro accuracy: | 83.0 % |

### Confusion matrix

In Table 3 the numeric column headings represent: 0: baboon, 1: buffalo; 2: bushbuck; 3: bushpig; 4: civet; 5: elephant; 6: genet; 7: hare; 8: hartebeest; 9: impala; 10: kudu; 11: nyala; 12: oribi; 13: porcupine; 14: reedbuck; 15: sable Antelope; 16: vervet Monkey; 17: warthog; 18: waterbuck; 19: wildebeest.

**Figure 7.**
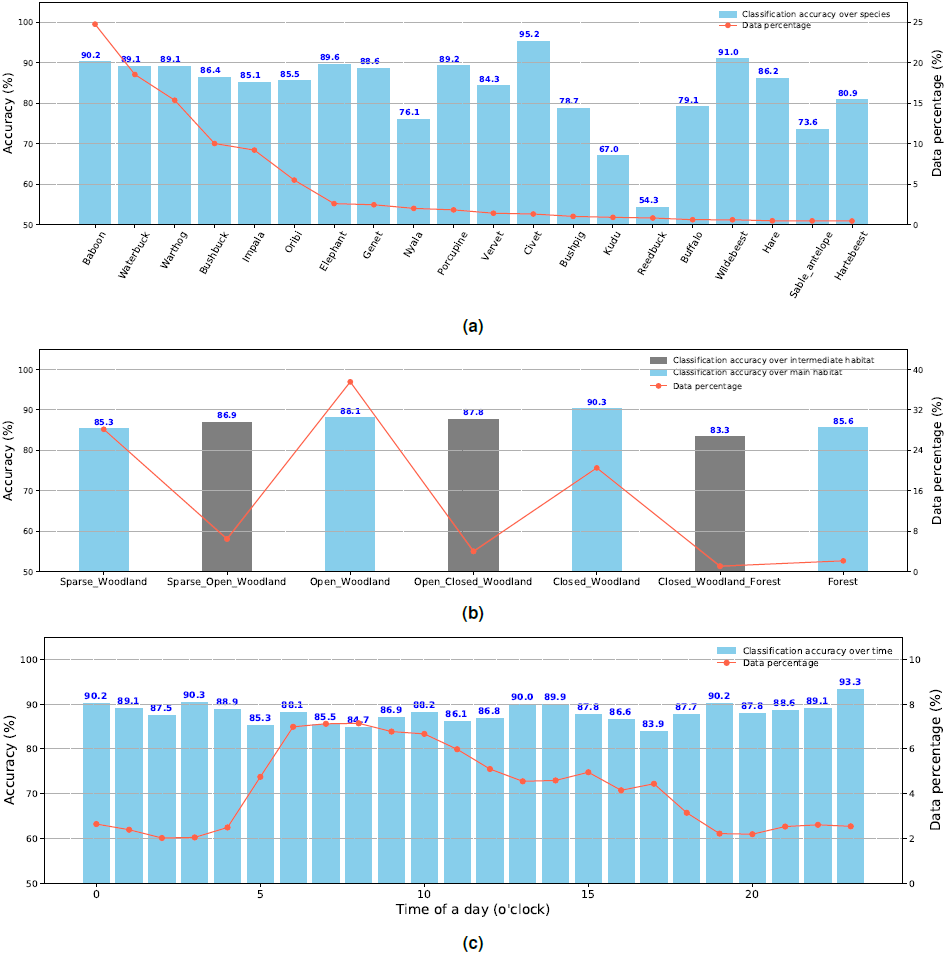
Model performance. (a) Animal classification accuracy and data distribution species. Reedbuck and Kudu are the two species for which the results were least accurate. (b) Animal classification accuracy and distribution by habitat type, (c) Animal classification and distribution by time of day. Unlike per-class accuracy, habitat and time accuracy are more evenly distributed, despite the overall data distribution. Class-aware sampling weakens the relationship between data distribution and accuracy so that the accuracy of data at the tail end of the distribution does not drop drastically.

**Table 3.** Confusion matrix in percentage (%)

|  | 0 | 1 | 2 | 3 | 4 | 5 | 6 | 7 | 8 | 9 | 10 | 11 | 12 | 13 | 14 | 15 | 16 | 17 | 18 | 19 |
| --- | --- | --- | --- | --- | --- | --- | --- | --- | --- | --- | --- | --- | --- | --- | --- | --- | --- | --- | --- | --- |
| 0 | 90.2 | 0.2 | 0.5 | 0.1 | 0.0 | 0.0 | 0.0 | 0.0 | 0.0 | 1.5 | 0.0 | 0.8 | 0.6 | 0.0 | 0.1 | 0.0 | 0.4 | 2.9 | 2.3 | 0.0 |
| 1 | 4.7 | 79.1 | 2.0 | 0.0 | 0.0 | 4.7 | 0.0 | 0.0 | 0.7 | 0.0 | 0.0 | 0.0 | 0.0 | 0.7 | 0.0 | 2.7 | 0.0 | 0.7 | 4.1 | 0.7 |
| 2 | 2.6 | 0.0 | 86.5 | 0.2 | 0.3 | 0.4 | 0.5 | 0.6 | 0.0 | 1.6 | 0.3 | 0.8 | 1.5 | 0.5 | 0.4 | 0.0 | 0.0 | 1.2 | 2.5 | 0.0 |
| 3 | 2.9 | 0.0 | 5.3 | 78.7 | 0.0 | 0.0 | 0.8 | 0.0 | 0.0 | 0.8 | 0.4 | 0.0 | 2.0 | 1.6 | 0.0 | 0.0 | 0.0 | 4.1 | 2.5 | 0.4 |
| 4 | 0.7 | 0.0 | 1.1 | 1.1 | 95.2 | 0.0 | 0.4 | 0.0 | 0.0 | 0.0 | 0.0 | 0.0 | 0.0 | 0.0 | 0.0 | 0.0 | 0.0 | 1.5 | 0.0 | 0.0 |
| 5 | 2.6 | 0.0 | 1.3 | 0.2 | 0.0 | 89.6 | 0.2 | 0.3 | 0.0 | 0.2 | 0.0 | 0.0 | 0.0 | 0.3 | 0.0 | 0.3 | 0.0 | 3.0 | 1.8 | 0.2 |
| 6 | 0.0 | 0.0 | 2.9 | 0.6 | 1.0 | 0.0 | 88.6 | 1.1 | 0.0 | 0.4 | 0.0 | 0.2 | 1.3 | 1.5 | 0.4 | 0.2 | 0.0 | 1.0 | 1.0 | 0.0 |
| 7 | 0.0 | 0.0 | 5.7 | 0.0 | 0.0 | 0.0 | 2.3 | 86.2 | 0.0 | 0.0 | 0.0 | 0.0 | 2.3 | 1.1 | 0.0 | 0.0 | 0.0 | 0.0 | 2.3 | 0.0 |
| 8 | 1.1 | 1.1 | 1.1 | 0.0 | 0.0 | 0.0 | 0.0 | 0.0 | 80.9 | 4.5 | 1.1 | 2.2 | 0.0 | 0.0 | 1.1 | 0.0 | 0.0 | 0.0 | 6.7 | 0.0 |
| 9 | 3.7 | 0.0 | 1.4 | 0.2 | 0.0 | 0.1 | 0.0 | 0.0 | 0.0 | 85.1 | 0.1 | 0.6 | 2.1 | 0.0 | 0.5 | 0.0 | 0.7 | 1.2 | 4.1 | 0.1 |
| 10 | 9.2 | 0.0 | 1.1 | 1.1 | 0.0 | 0.0 | 0.0 | 0.0 | 0.0 | 3.2 | 67.0 | 3.8 | 1.1 | 0.0 | 0.0 | 0.0 | 0.0 | 3.8 | 9.2 | 0.5 |
| 11 | 4.1 | 0.0 | 6.9 | 0.0 | 0.0 | 0.5 | 0.0 | 0.0 | 0.0 | 3.9 | 1.4 | 76.0 | 0.0 | 0.2 | 0.0 | 0.5 | 0.2 | 0.2 | 5.8 | 0.2 |
| 12 | 2.4 | 0.0 | 3.4 | 0.0 | 0.0 | 0.0 | 0.3 | 0.2 | 0.2 | 2.6 | 0.3 | 0.0 | 85.5 | 0.2 | 0.5 | 0.0 | 0.0 | 1.7 | 2.8 | 0.0 |
| 13 | 0.8 | 0.0 | 3.5 | 1.1 | 0.0 | 0.3 | 1.9 | 0.5 | 0.0 | 0.8 | 0.0 | 0.0 | 0.0 | 89.3 | 0.0 | 0.0 | 0.0 | 0.5 | 1.3 | 0.0 |
| 14 | 1.1 | 0.0 | 11.4 | 0.0 | 0.0 | 0.0 | 1.7 | 0.0 | 0.0 | 11.4 | 0.6 | 0.6 | 8.0 | 0.0 | 54.3 | 0.0 | 0.0 | 0.6 | 10.3 | 0.0 |
| 15 | 1.7 | 3.3 | 3.3 | 0.8 | 0.0 | 0.8 | 0.0 | 0.0 | 1.7 | 1.7 | 0.8 | 0.8 | 0.0 | 0.0 | 0.0 | 73.6 | 0.0 | 1.7 | 9.9 | 0.0 |
| 16 | 8.0 | 0.0 | 0.6 | 0.0 | 0.0 | 0.0 | 0.0 | 0.0 | 0.0 | 1.5 | 0.0 | 0.9 | 0.3 | 0.0 | 0.0 | 0.0 | 84.3 | 1.2 | 3.1 | 0.0 |
| 17 | 4.8 | 0.1 | 0.9 | 0.2 | 0.1 | 0.3 | 0.0 | 0.0 | 0.0 | 0.9 | 0.0 | 0.2 | 0.8 | 0.0 | 0.1 | 0.0 | 0.4 | 89.1 | 1.9 | 0.1 |
| 18 | 2.3 | 0.1 | 1.2 | 0.0 | 0.0 | 0.2 | 0.0 | 0.1 | 0.0 | 2.0 | 0.1 | 0.1 | 0.5 | 0.0 | 0.2 | 0.1 | 0.4 | 3.2 | 89.1 | 0.1 |
| 19 | 0.0 | 0.0 | 0.0 | 0.0 | 0.0 | 0.0 | 0.0 | 0.0 | 2.7 | 0.0 | 0.0 | 0.0 | 0.0 | 0.0 | 0.0 | 0.9 | 0.0 | 0.9 | 4.5 | 91.0 |

## Appendix 4: Interpretation methods details

### Outline of our general approach

**Figure 8.**
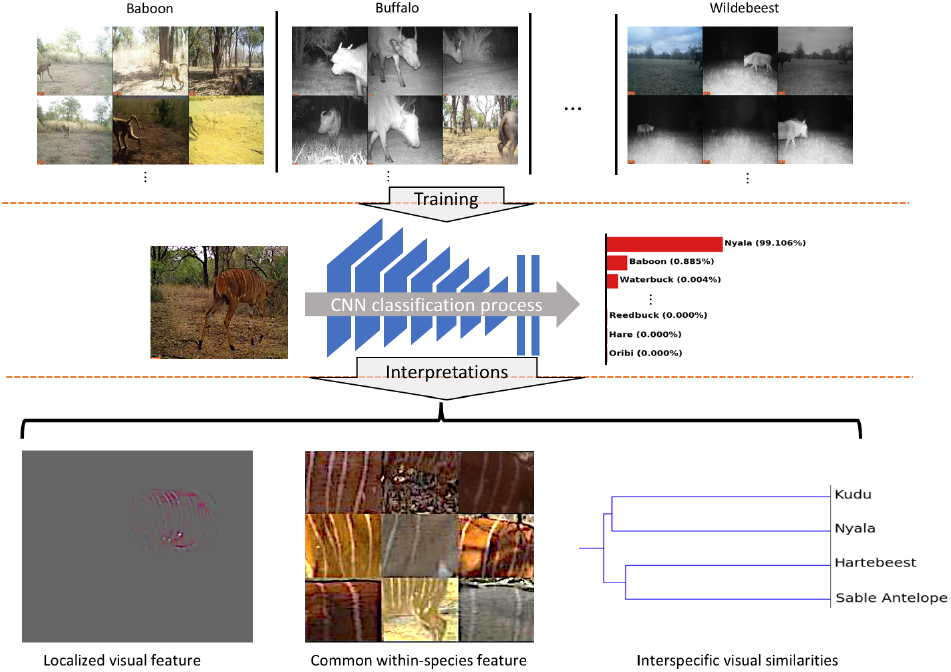
Overview of training and interpretations. We use the VGG-16 algorithm to train a convolutional neural network (CNN) on a camera-trap dataset collected from Gorongosa National Park, Mozambique. When a trained CNN is fed with images, it classifies the images with a classification probability. To interpret the mechanisms behind this process, we use Guided Grad-CAM (GG-CAM) to find the localized visual features extracted by the CNN from the images. Next, we use Mutual Information (MI) on the neurons of the last convolutional layer to generate common within-species features of each species. Finally, we use hierarchical clustering on the feature vectors to study the relative interspecific visual similarities of each species in the dataset. Comparing CNN with human knowledge, we find that the features generated by a trained CNN for animal identification are similar to those used by humans.

### Guided Grad-CAM (GG-CAM)

GG-CAM is a method that combines the output of Grad-CAM and GBP^14^:

Grad-CAM generates coarse, discriminative regions according to animal species. It is calculated as the rectified linear units (i.e., that is the function max*{*0,*x}*) of the weighted sum of the response maps from the last convolutional layer (Eq. 1). The weighted sum is based on the importance value *α*_*k*_ (importance value of the *k_th_* neuron) of each neuron (neuron importance) in that layer of the response map, *A*^*k*^, where its *i j*^*th*^ element is 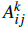 for a total number of elements *Z*. If *y* is the prediction score of animal A before the Softmax layer, then GG-CAM is computed using the following equations:

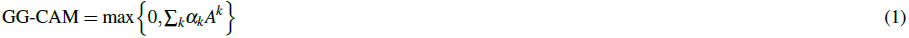

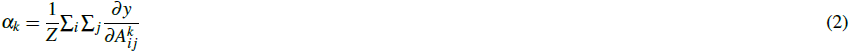

GBP is a method that captures non class-discriminative details of visual components that are important to the network overall. It is calculated as the gradient of the output response map of the last convolutional layer with respect to the input image, with only positive gradients and positive response elements (Eq. 3). As in^14^, we then combine the products of Grad-CAM and GBP as the final results of animal discriminative features in each image. Figure 9 depicts the difference between Grad-CAM, GBP, and GG-CAM (the combination of the former two methods). If *R*^*l*^ is the GBP product of the *l*^*th*^ layer then it is calculated in terms of the response maps *f*^*l*^ of the *l*^*th*^ layer, and the response maps *f*^out^ of the last convolutional layer. Specifically, defining 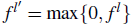, the equation is:

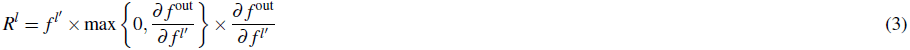

**Figure 9.**
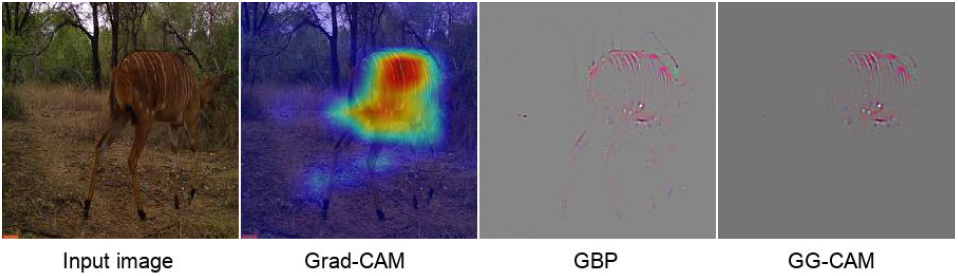
Comparison between Grad-CAM, GBP, and GG-CAM. Once trained, any image (leftmost panel) can be overlaid with its Grad-CAM heat map (left center panel) to identify the region of ‘most interest’ to the CNN (see Appendix 4). Similarly, the corresponding feature map (center right panel), produced using Guided Back-propagation (GBP), (which, as described in Appendix 4, identifies the most important visual features to our CNN) can be weighted by the Grad-CAM heat map to produce the guided Grad-CAM (GG-CAM) image seen in the rightmost panel. Note that in this Nyala image, GBP is less discriminative than GG-CAM: both highlight the stripes of the Nyala, whereas GPB includes non-species-discriminative tree branches and legs.

### Mutual information

Next, we demonstrate another approach to inspect within-species animal discriminative features based on common neuron importance. Each neuron in the network has a response to certain parts of the input images. Classification of images is based on a combination of the neuron responses. In addition, per species, certain neurons are more important than others for classification. We assume that the responses from these neurons can be regarded as common within-species features.

We use Mutual Information (MI)^16^, a method commonly used to find information shared between variables^15, 48^, on the neuron importance (Eq. 2) (normalized from 0 to 1) from the last convolutional layer across the data (Eq. 4). The results are illustrated in Figure 11 for image patches with the highest responses to the neurons with the top 1 to top 5 mutual information scores of each species.

We calculated *I*(*U* < *C*), the MI for neuron *U* and animal species *C*, as follows. Suppose *N*_11_ and *N*_01_ are the number of images of *C*, where *U* has neuron importance > 0.5 and ≤ 0.5 respectively. Further suppose *N*_10_ and *N*_00_ are the number of images that are not C, where U has neuron importance > 0.5 and ≤ 0.5 respectively. Defining *N*_1_*_•_* = *N*_10_ +*N*_11_, *N_•_*_1_ = *N*_11_ +*N*_01_, *N*_0_*_•_* = *N*_00_ + *N*_01_, *N_•_*_0_ = *N*_00_ + *N*_10_, and *N* = *N*_00_ + *N*_01_ + *N*_10_ + *N*_11_ it then follows that

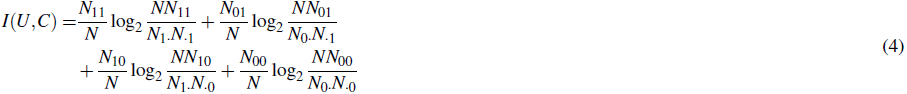

### Interspecific visual similarities and species familiarity

To inspect visual similarities between animal species, we generated a visual tree of all species by implementing hierarchical clustering on the feature vectors before the classifier layer. Firstly, we extracted the feature vectors of 6000 randomly selected training images and applied Principal Component Analysis (PCA) to compress the 4096-dimension feature vectors to 128 dimensions for computational simplicity. We then computed the average interspecific Euclidean distances between every pair of the 20 species. Finally, we processed the interspecific distances using a hierarchical clustering method with the Ward variance minimization algorithm^49^.

Similarly, the relative familiarity of 30 animals species was calculated as the Euclidean distances of each animal species within the tree. We used an experimental testing dataset (Data Section), which contains 20 randomly selected images of each species. We treated each of these 30 species as a new species (either known or unknown) and generated a new similarity tree based on the 30 “new” species and the 20 “old” species. Then, we calculated the Euclidean distances between each of the 30 “new” species and their closest “old” species. The larger the distance, the more unfamiliar the species is to the CNN.

## Appendix 5: Additional results

### Extracted feature similarities

**Figure 10.**
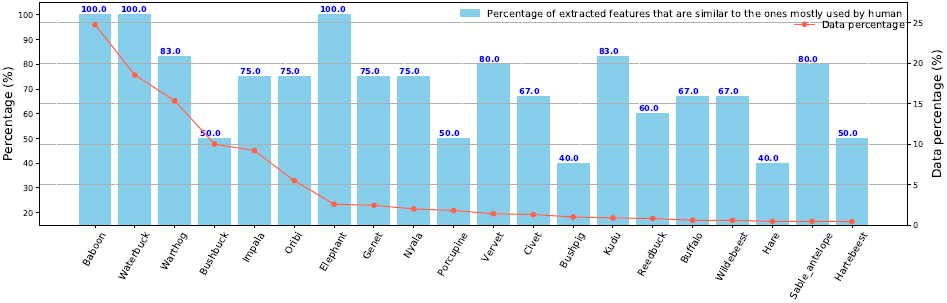
Percentage of extracted features that are similar to corresponding visual descriptors of each species created before these analyses started (Table 4). The similarity was agreed upon by up to four authors (ZM, KMG, ZL and MSN), who scored 9 randomly selected images for each species. The extracted features are mostly similar to or correspond to human descriptors. The higher the percentage, the more similar these features are to human visual descriptors.

### Mutual information results of all animal species

**Figure 11.**
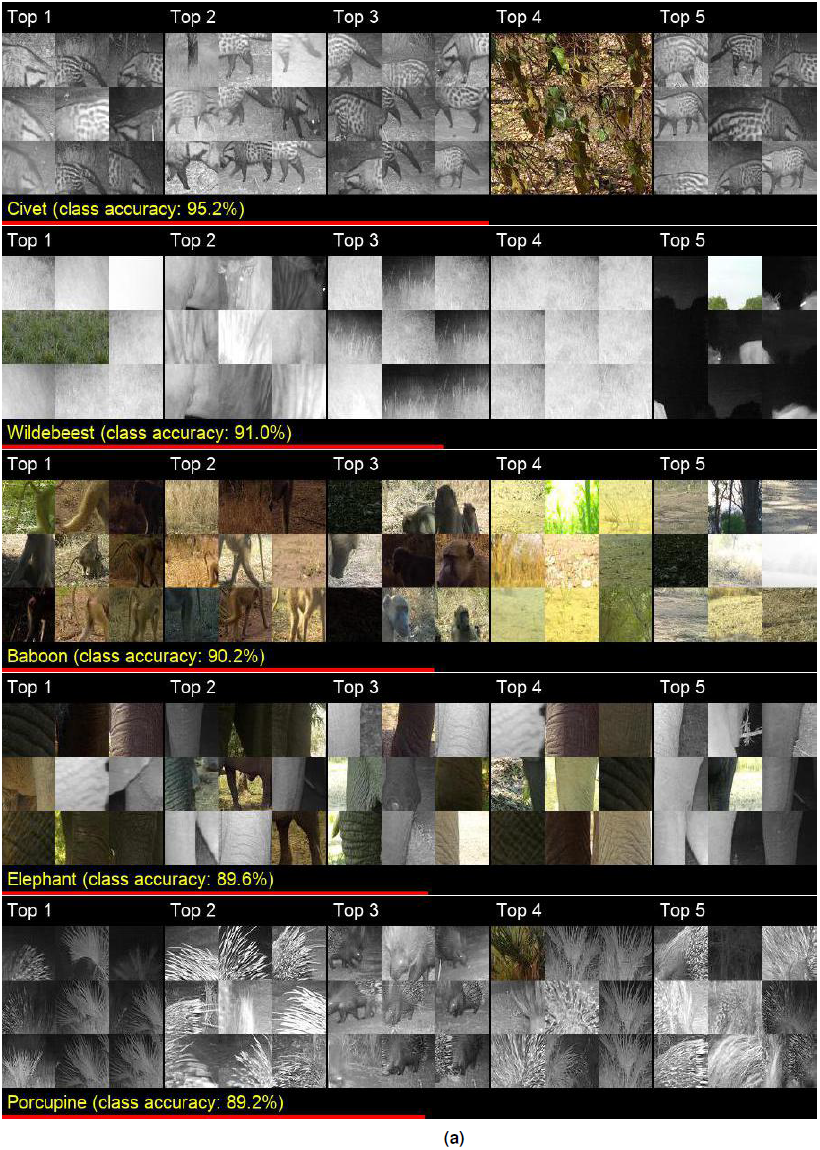
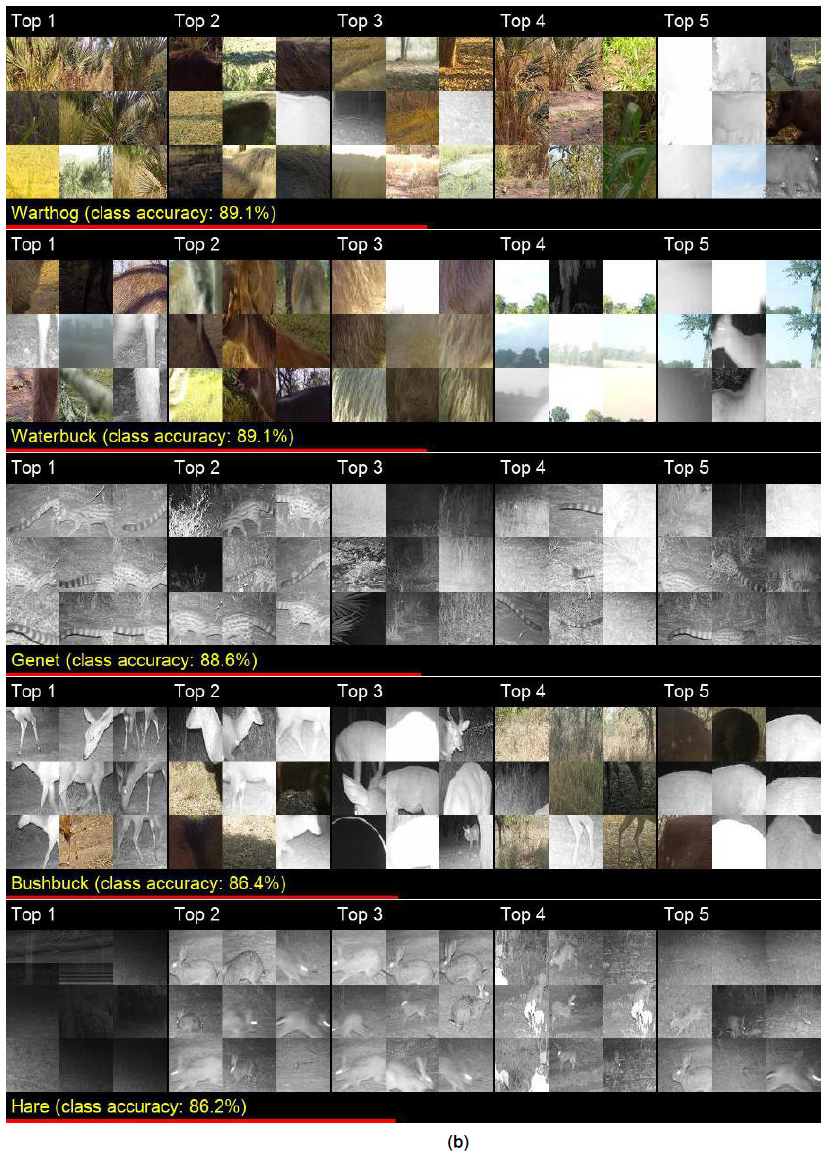
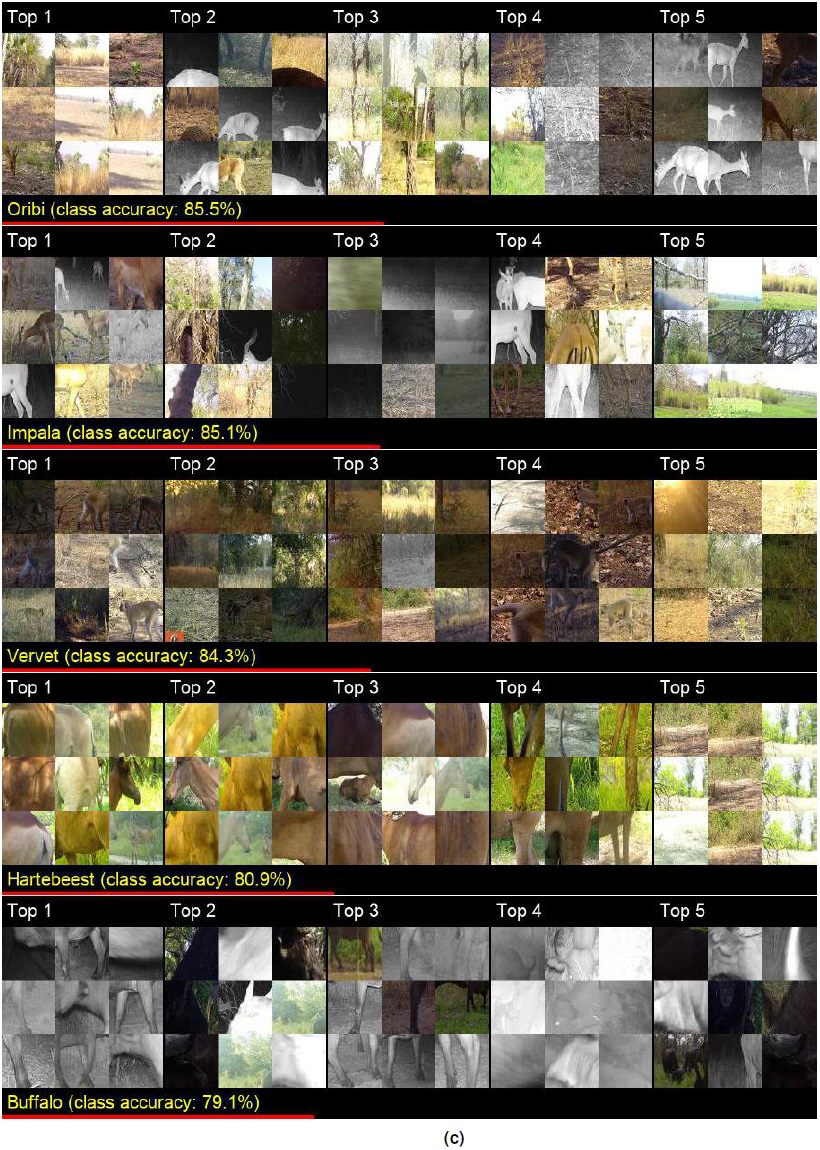
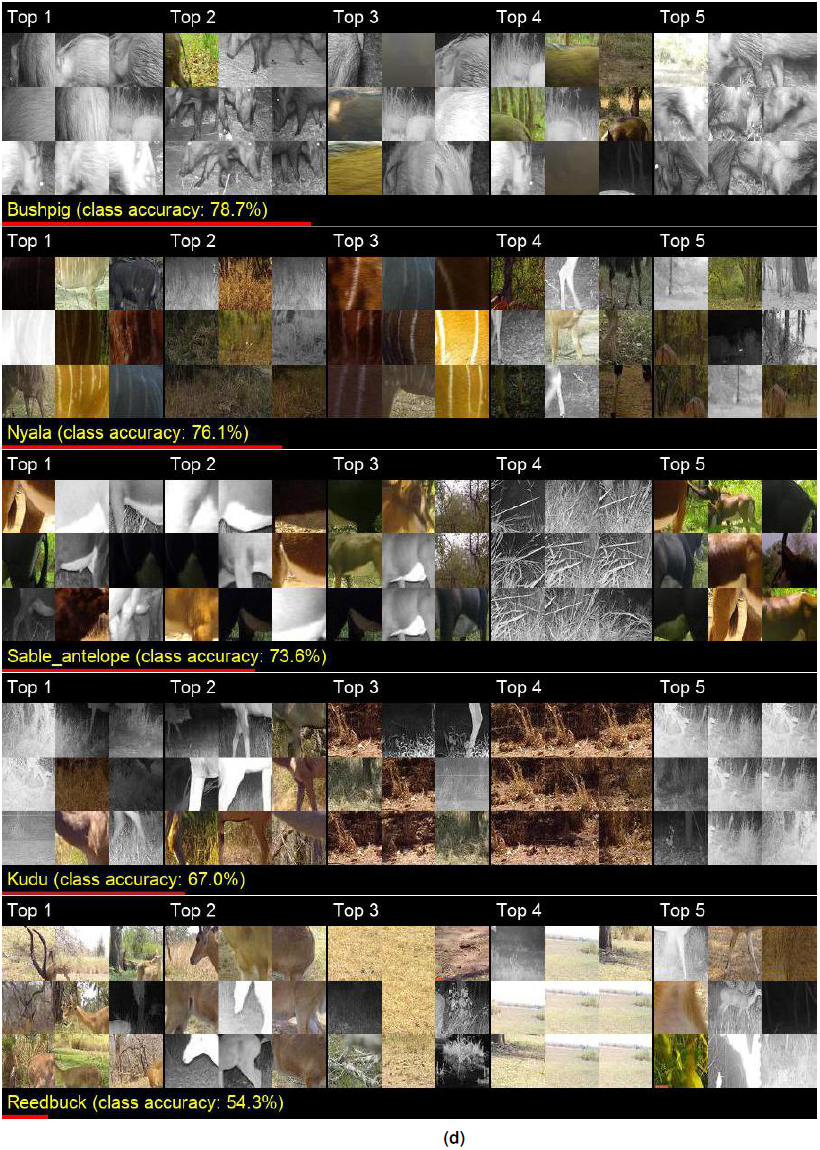
Following Figure 2 for the case of porcupine and reedbuck, here the extracted patches are centered around the hottest pixel of the five most responsive neurons in the last convolutional layer of our CNN that has the highest MI score (Methods) for all 20 species (i.e., the first two are repeated for the sake of completeness). Red bars are graphical representations of class accuracy.

## Appendix 6: Full visual descriptors

**Table 4.** Features used by humans to identify the 20 most common species from camera trap images. These features were identified through a survey of people with extensive experience in classifying camera trap data from Gorongosa. The features below were selected by at least 5 of the 13 survey respondents.

|  |  |
| --- | --- |
| **Baboon** | primate body type<br>tail curving upward at base<br>long, dark snout |
| **Buffalo** | horns that curve to the side of head<br>stocky barrel-shaped body<br>dark coat |
| **Bushbuck** | thick ring of short, dark fur along neck<br>parallel, slightly spiraled horns<br>rounded rump<br>ungulate body type<br>white spots along the rump |
| **Bushpig** | pig body type<br>silver-colored mane |
| **Civet** | nocturnal<br>small carnivore body type<br>black spots<br>rounded back<br>short, black legs<br>crest of black hair from head to tail |
| **Elephant** | stocky, rectangular body shape<br>gray to brownish wrinkled skin<br>long trunk<br>huge ears that are wide at base and narrow at bottom<br>thick, round legs<br>white tusks |
| **Genet** | slender body |
|  | long, narrow tail<br>banded tail<br>black spots<br>small carnivore body type |
| **Hare** | round body<br>long ears that point up |
| **Hartebeest** | curved horns<br>ungulate body type<br>uniform dark brown coat |
| **Impala** | S-shaped horns of male<br>tri-colored body<br>black streaks on the rear |
| **Kudu** | long horns with large spirals (males)<br>hump on back of neck<br>thin, white stripes on back<br>white band between the eyes<br>light gray/brown color<br>ungulate body type<br>long, slender legs |
| **Nyala** | thin, white stripes on back<br>golden fur of female, dark brown fur of male<br>white spots on face and nose of male<br>spiral horns of male<br>ungulate body<br>white and yellow leg markings |
| **Oribi** | short, straight horns of male<br>white abdomen<br>short, black tail<br>black circular patches under ears<br>conical head shape<br>ungulate body type |
| **Porcupine** | nocturnal |
|  | long black and white quills<br>stout, rounded body shape |
| **Reedbuck** | forward-curving horns of male<br>black circular patches under ears<br>uniform coloration<br>ungulate body shape |
| **Sable antelope** | long, backward-curving horns<br>horse-like body type<br>white striped facial markings<br>white underbelly<br>chestnut coat of female and dark brown color of male |
| **Vervet** | primate body type<br>black face<br>long tail, held out straight<br>white brow |
| **Warthog** | pig body type<br>two pairs of upward-pointing tusks<br>mane from top of head to middle of back<br>thin tail with tuft of hair at the bottom<br>flat, wide snout |
| **Waterbuck** | ribbed horns, curved out and forward (male)<br>white circular ring of fur on rump<br>shaggy, coarse, red-brown fur<br>black nose<br>ungulate body type |
| **Wildebeest** | curved horns that are wider than they are tall<br>horse-like body type<br>long, rectangular face<br>black beard<br>black mane along back<br>black vertical stripes on neck |

